# Anticipating others’ future behaviours alters collective movement and enhances survival

**DOI:** 10.1101/2025.08.14.670290

**Authors:** Pranav Minasandra

## Abstract

Current models of collective animal movement explain several emergent phenomena. However, these models do not account for animals’ potential ability to forecast each others’ decision-making processes. The ability to predict and account for the behaviours of others is likely unique to animals, and the relevance of this ability in collective movement merits exploration. I granted anticipatory abilities to Hamilton’s selfish herd, a classic model of collective behaviour, using 14000 simulated populations and explored the consequences of this ability on collective movement patterns. I also simulated competitions between agents with and without anticipatory abilities, and showed that anticipation may confer a survival advantage.

## Introduction

Social animals make collective movement decisions. Most models of collective movement rely on spatial interaction rules, exploring various decentralised emergent phenomena [1–3]. Such models feature relatively simple interaction rules such as attraction, alignment, and repulsion that allow individuals to constantly tune their position and velocity. These models have helped explain and elucidate many real-world patterns observed across species [4–6]. However, real animals can be cognitively sophisticated, and their movement is often guided by more complex rules than instantaneous attraction and repulsion [e.g., 3, 7].

A sufficiently cognitively advanced animal may not only react to the current and remembered past states of other animals, but also consider their forecasted future behaviours. In collectives, anticipation could help individuals avoid collisions [8] and conflicts, and improve coordination in cooperative tasks [9]. Most anticipation models are physics-based, and perform linear extrapolations of individual’s trajectories [10, 11]. However, anticipation may also stem from cognitive abilities like the theory of mind [12, 13], wherein animals can attest mental states to each other. Cognitively sophisticated animals may mentally model other individuals’ decision-making processes. With this ability, animals may anticipate each others’ behaviours using embedded mental models (which I will call *metarepresentations*) of each others’ decision-making processes. Metarepresentations would allow animals to incorporate their conspecifics’ perception of social and environmental variables, leading to better predictions of their movement decisions than mere extrapolation of trajectories. Despite their relevance in social interactions, extrapolative anticipation has received limited attention, and metarepresentational attention practically none, in modelling collective movement. Here, I augment a classic model of collective movement, Hamilton’s Selfish Herd [14] with metarepresentational anticipation. I then quantify the impact of anticipation on collective movement and survival probabilities.

## Modelling anticipation

To explore the effect of anticipation on collective movement, I performed simulations where I granted varying levels of metarepresentational anticipation to populations of different sizes. I started with agents moving according to a baseline movement model, which I called *d*_0_ (see next paragraph). Next, I allowed these fundamentally naïve agents to anticipate each others’ movement decisions and update their own decisions appropriately. When an individual could anticipate others’ behaviours, it predicted everyone’s movements according to the *d*_0_ model, and updated its movement decision with respect to these predicted future locations of the others. I called this new model where all individuals move with updated decisions, *d*_1_. However, all agents may further use metarepresentational anticipation to predict others’ *d*_1_ positions, and before moving, perform a second update, which yields the *d*_2_ model of movement. Another recursion in anticipation similarly yields the *d*_3_ model, and this logic can be applied *ad infinitum*. Starting from a baseline model, I thus simulated three additional movement models featuring increasing anticipatory abilities (Figure 1).

**Figure 1:**
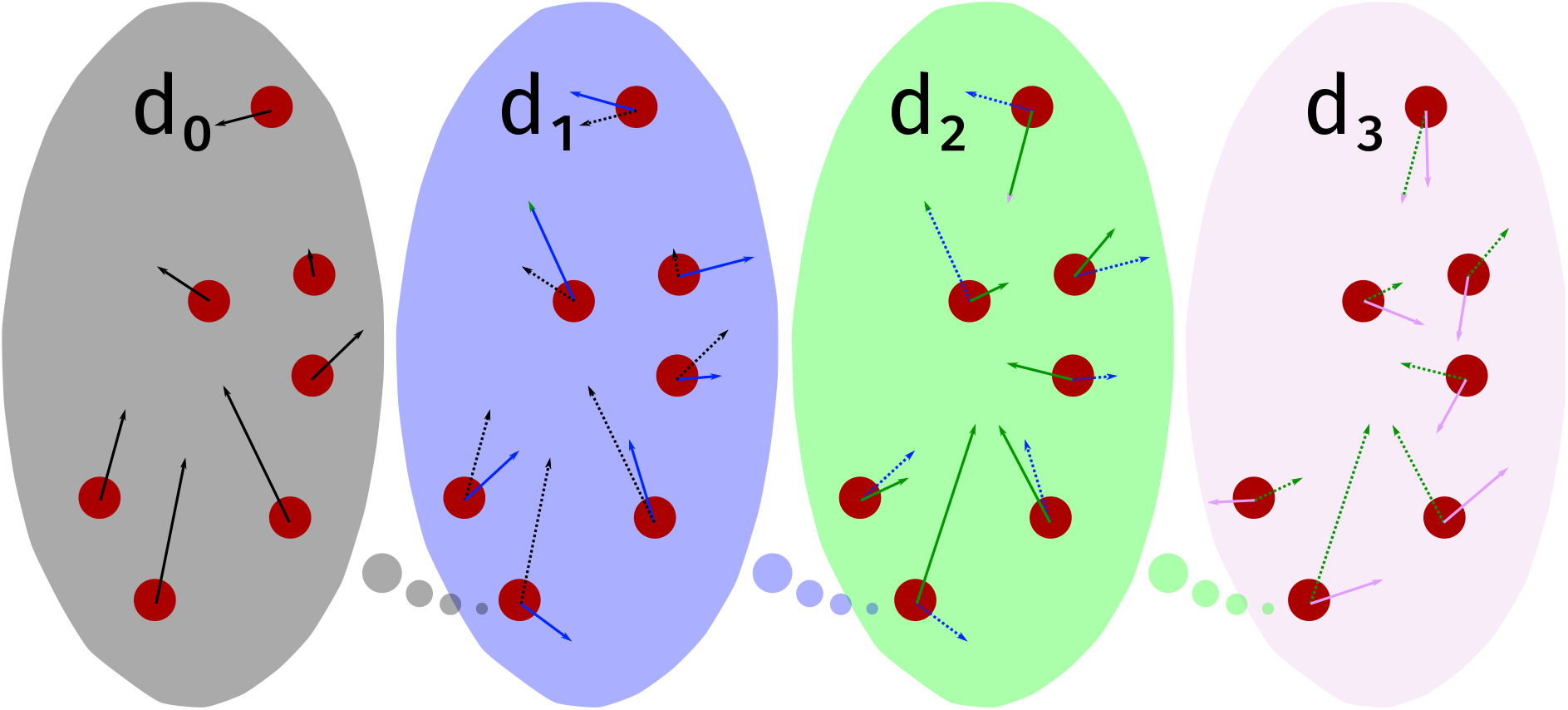
Recursively updating movement rules with anticipation of others’ decisions leads to an infinite sequence of models. *d*_0_ individuals move according to a baseline model (black arrows) and do not anticipate others’ behaviours. *d*_1_ individuals predict others’ *d*_0_ movement decisions, and appropriately update their movement rules (blue arrows). This way, individuals that mentally emulate the *d*_*j™*1_ movement model can update their decisions based on these predictions, yielding the *d*_*j*_ movement model.

As *d*_0_ I chose Hamilton’s Selfish Herd, a biologically motivated collective behaviour model of group formation. [14] Each agent made constant movement decisions to minimise the area of its *domain of danger*, the area around the agent in which predators would likely choose that agent as their exclusive target. Per Hamilton, an agent’s domain of danger could be proxied by its Voronoi polygon [14], and the area of this polygon captured the animal’s predation risk. A domain of danger minimisation movement rule leads to individuals, driven by the emergent push to occupy locations between other individuals, aggregating into groups; and to some individuals coming to rest eventually on the edge of the simulation arena. With anticipation, agents were programmed to minimise their domains of danger considering the predicted locations of all others.

## Results

How did models with increased depths of anticipation (*d*_1_, *d*_2_ and *d*_3_) differ from the baseline selfish herd? I initialised 500 populations of each population size, and let these agents move according to each of *d*_0_—*d*_3_ for 500 iterations. This was done for seven population sizes.

### More polarised group movement

To characterise group movement, I quantified the polarisation of groups, i.e., the degree to which individuals were moving in similar directions. To incorporate only freely moving groups, I excluded groups on the arena edge. The median polarisation of groups, at all population sizes, increased sharply with the depth of anticipation.(Figure 2A)

**Figure 2:**
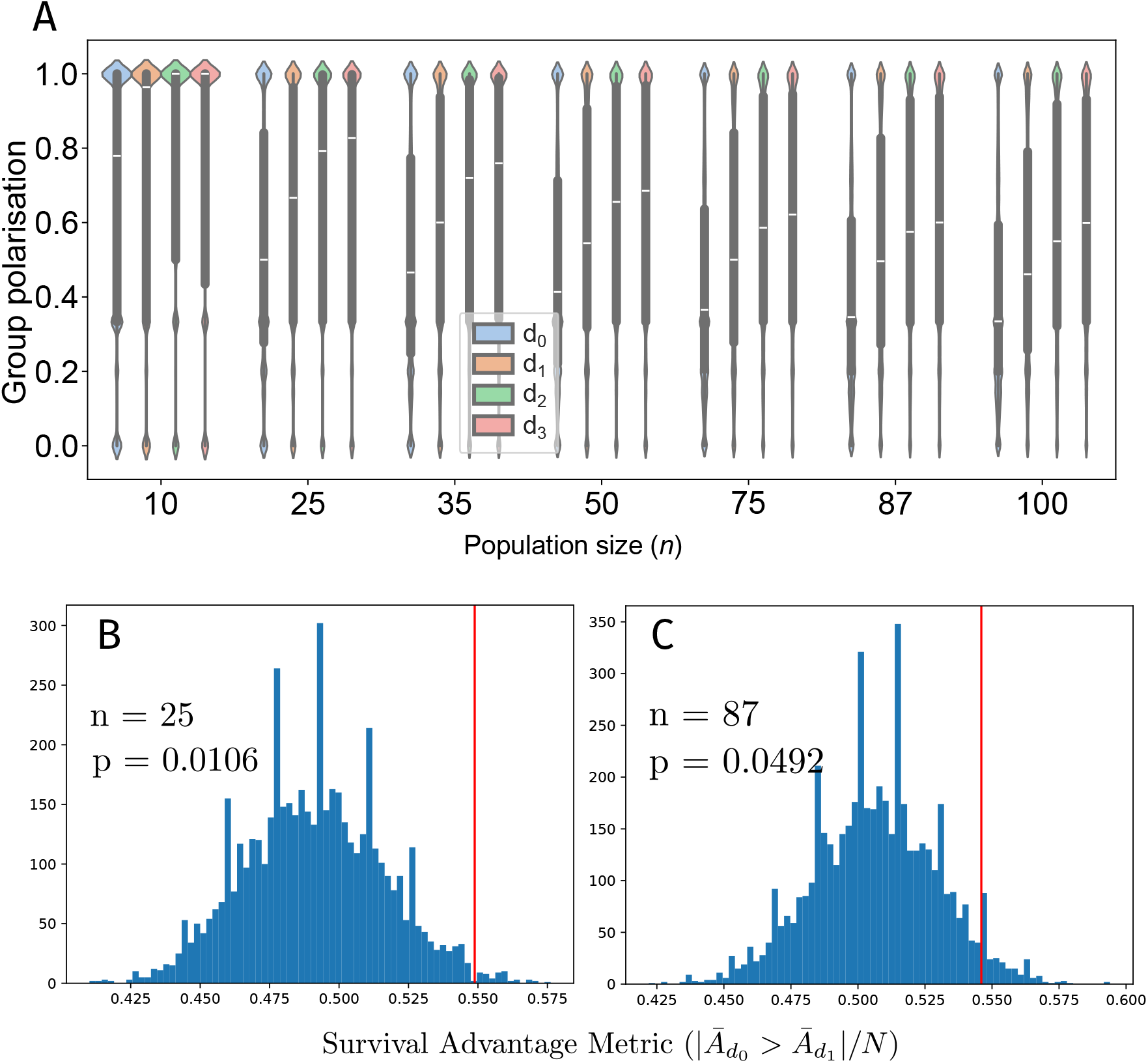
(A) With increasing depth of anticipation, groups show increased median polarisation. For each population size I show four violins for *d*_0_—*d*_3_. Small white lines are medians, while gray rounded rectangles show 0.25—0.75 interquantile intervals. (B, C) Metarepresentational anticipation reduces the size of domains of danger in a competition trial in 2 population sizes. Red lines show the true value of the survival advantage metric, while histograms in blue show this metric’s distribution across individuals (see Methods).

### An evolutionarily stable strategy

Can metarepresentational anticipation alone can provide an evolutionary advantage? I simulated two *d*_0_ populations (*n* = 25 and 87) each invaded by five *d*_1_ individuals. I predicted that anticipation would enable *d*_1_ individuals to occupy central locations in groups, and thereby lessen their predation risk.

Each population was initialised with 500 randomised locations and each initialisation run for 500 iterations. For *d*_1_ individuals, I then computed a survival advantage metric (the probability of *d*_1_ individuals having a smaller domain of danger on average; see Methods), and compared this value to the null distribution of the same metric. I found that in each population, the mean area of the *d*_1_ individuals’ domains of danger was significantly smaller than expected from chance alone (*p* = 0.011 and *p* = 0.049 respectively, Figure 2B and C)

## Discussion

I provided a way to generate a sequence of augmented movement models considering individuals that anticipate each others’ behaviours using metarepresentations of decision-making processes. Applying this method to Hamilton’s Selfish Herd, I have shown the emergence of a polarised collective movement pattern in groups and a competitive advantage experienced by anticipation-capable individuals over others in a naïve population.

The movement model changed qualitatively and quantitatively with anticipation. Groups moved with increased polarisation when individuals could anticipate each others’ behaviours. Selfish herd members typically seek to occupy central locations within groups, but without anticipation, individuals’ uncoordinated decisions may hamper the effective reduction of their domains of danger. Anticipation may enable individuals to avoid repeated futile attempts to reach the group centre, thereby improving group coordination. Anticipation-capable individuals also experienced smaller domains of danger in mixed *d*_0_-*d*_1_ populations, and therefore lower predation risks, compared to naïve individuals. This survival advantage conferred by anticipation, even if small, may lead to its evolutionary stability.

The approach I used to augment Hamilton’s model here can be applied readily to other movement models, particularly by rewriting them as continuous optimisations of some potential function. Extrapolative anticipation has been expected to have a stabilising effect [10], and I suspect that the increased polarisation (or in a sense, ‘consensus’) of groups capable of metarepresentational anticipation is analogous to this effect. It is worth examining whether such a stabilising effect, or another generalisable result, manifests across movement models augmented with anticipation.

When an individual constructs a metarepresentation of another, it must also account for that individual’s own metarepresentations, resulting in recursive complex cognitive exercises involving nested metarepresentations requiring significant cognitive infrastructure and effort. However, decision-making rules that merely approximate metarepresentational anticipation could arise from animals responding to behavioural cues, and could function more as intuition than as explicitly simulated mental models. Moreover, with metarepresentational anticipation, incorrect assumptions in lower layers of anticipation can be subsequently magnified leading to suboptimal decisions, leading to the possibility of ‘overthinking’ decisions. Strategies that optimally combine *d*_0_ and *d*_*j*_ decisions might lessen this effect. Incomplete and/or imperfect information during anticipation also thus merit further theoretical study.

In a setting free from costs, confounds, and cognitive constraints, I have shown that metarepresentational anticipation, the ability of animals to explicitly emulate others’ thought processes to predict future behaviours, can play a large role in collective movement. Cognitive capacities and abilities like theory of mind vary substantially across the tree of life [15], and understanding the extent and nature of this effect may be complicated. Understanding the role of anticipation in real-world systems is a challenging problem with interdisciplinary ramifications. A better understanding can be accomplished from further simulation studies as well as novel analyses using field data.

## Methods

(Additional details about methods in Extended methods.)

### Simulations

The selfish herd was simulated by computing the areas of the Voronoi polygons of all agents and executing gradient descent algorithms by each agent on its Polygon’s area. Agents were confined to the unit square, whose boundaries also limited the Voronoi polygons. For population sizes *n* = 10, 25, 35, 50, 75, 87, and 100, I ran *d*_0_ · · · *d*_3_ models 500 times for 500 iterations each. The *d*_*j*_th movement model worked by running the *d*_*j−*1_th model each iteration, then returning each individual to its position in the previous iteration and allowing it an additional round of gradient descent to minimise the size of its predicted Voronoi polygon.

### Metrics

From these trajectories, at *t* = 40, 60, …, 200, I computed the polarisation of each group. Polarisation was computed as 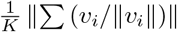, where *v*_*i*_ is the velocity vector of the *i*th individual and *K* is the group size. Groups at the edge of the arena were excluded.

### Competition trials

I simulated 500 populations with five *d*_1_ individuals and all others at *d*_0_, *n* = 25 and 87. Populations were iterated 500 times. At *t* = 250, 270, …, 350, I computed the mean areas of the Voronoi polygons for all *d*_1_ individuals,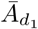. I computed a survival advantage metric: the proportion of simulations where the 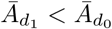. To test for statistical significance, I randomly chose five *d*_0_ individuals from the population 5000 times, to compute the null distribution of this metric, assuming the randomness arose solely from finite-size effects. The true value was compared to this null distribution to obtain the *p*-value with one tail.

## Acknowledgements

I thank Cecilia Baldoni for her very useful comments during the course of this study. I also thank Alison Ashbury, Gabriella Gall, Carel van Schaik, Ariana Strandburg-Peshkin, and Raghavendra Gadagkar for valuable discussions. When doing part of this work, I was funded by an IMPRS-QBEE Postdoctoral Award.

## Code availability

All code for these simulations and analyses can be found on GitHub: github.com/pminasandra/selfish-herd-social- competence.

## Appendix A Extended methods

### Setup

The populations of agents I simulated followed the same rules proposed by Hamilton in his classic selfish herd theory paper [14]. At every iteration, each agent computed its ‘domain of danger’. As per Hamilton’s formulation [14], an animal’s domain of danger was the set of all points in 2-dimensional space where said animal is closer than all other animals. This is also the animal’s Voronoi polygon. Each animal then moves in a way as to minimise its own domain of danger.

Here, I granted varying extents of anticipation to all agents in the selfish herd. I called these extents of anticipation *d*_0_, *d*_1_, *d*_2_, and *d*_3_. *d*_0_ individuals performed straightforward selfish herd movement with no anticipation of others’ movements. *d*_1_ individuals anticipated each others’ *d*_0_-model decisions, predicted where they would be in the next iteration, and updated their decisions so that the areas of their domains of danger would be minimised in that next iteration. *d*_2_ individuals accounted for each others’ *d*_1_ level updates to further refine their decisions, and *d*_3_ individuals performed this recursion one more time. It is technically possible to recurse indefinitely over these depths of reasoning *d*_*j*_, but I restricted ourselves to *d*_3_ because of explosively increasing computational costs. All simulations described here required 1 month to complete on an advanced computer (1 TB RAM, 40 threads).

I explored the collective movement of these anticipation-capable agents at seven different population sizes spanning an order of magnitude: 10, 25, 35, 50, 75, 87, and 100. For each population size, we chose 500 initialisation conditions (i.e., the initial locations of all agents). We then proceeded to perform 500 iterations each for *d*_0_, …, *d*_3_ starting from each of these conditions. Thus, a total of 7 *×* 500 *×* 4 = 14000 simulations of 500 iterations each were performed.

#### Calculation of domains of danger and movement rules

Following Hamilton’s characterisation, I assumed that each agent’s domain of danger was its Voronoi polygon. All calculations of Voronoi tessellations were done using the python library scipy (v1.15) [16]. To avoid Voronoi polygons with infinite areas, I restricted the agents to the unit square. For this restriction, I created dummy agents by mirroring the real agents about the edges of the unit square, and performed the calculation of Voronoi polygons including both real and dummy agents. I could then compute the area of the Voronoi polygon of any agent using the shoelace formula,

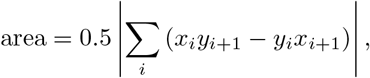

where (*x*_*i*_, *y*_*i*_) were all the vertices of its Voronoi polygon. By construction, the sum of Voronoi areas of all real agents was 1.0.

Naïve (*d*_0_) agents performed straightforward selfish herd movement. *d*_0_ agents moved by performing a gradient descent in 2D space to minimise their Voronoi areas. In each iteration, if an agent was located at (*x, y*) with a Voronoi area *A*(*x, y*), it sampled its new Voronoi areas 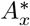 if it were to move to (*x* + *ε, y*) and 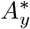 if it were to move to (*x, y* + *ε*). The gradient was then computed as

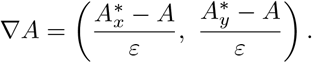

The agent then moved by performing a standard gradient descent update, (*x*^*\**^, *y*^*\**^) = (*x, y*) *−η ∇A*. In my simulations, I used *ε* = 0.005 and *η* = 0.1.

Anticipation was implemented in all *d*_*j>*0_ simulations by assuming that each agent was capable of mentally emulating the movement rules of *d*_*j−*1_ populations. Each *d*_*j*_ individual predicted the movement of the rest of the population based on *d*_*j−*1_ dynamics and computed the locations of all its conspecifics in the next iteration. Then, considering the forecasted positions of all others and the original position of itself, it computed the new location that minimised its Voronoi area in the next instant of time. *d*_1_, *d*_2_, and *d*_3_ dynamics were carried out using this recursive approach.

In all cases, the movement speed was capped so that each agent could not move more than 0.05 units per iteration.

#### Quantification of relevant metrics

After every twenty iterations, I extracted biologically relevant metrics pertaining to the positions of the animals. First, we quantified each animal’s predation risk by extracting the areas of each animal’s Voronoi polygon. I also used the clustering algorithm DBSCAN (scikit-learn v1.6.1) [17] with the parameter *ϵ* = 0.005 to label which group each animal belonged to [as in 18]. Lone animals were assumed to be groups of size 1.

I quantified the polarisation of a group as

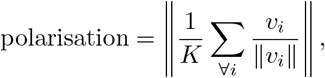

where *v*_*i*_ is the velocity vector for the *i*th individual in the group in that iteration, *K* is the group size, and *∥*… *∥* is the *L*^2^ norm. I computed this metric every 20th iteration between *t* = 40 and 200 for all groups with more than 1 individual. The restriction to *t* = 40 to 200 was to ensure that (a) effects of initial randomness were eliminated, and (b) a majority of groups had not migrated to the arena edge. Even within this time interval, groups with members on the edge of the arena were not considered, as their movement was restricted by the wall.

#### Competition simulations and analyses

I intended to measure differences in Voronoi areas in mixtures of agents with and without theory of mind. I considered populations of 25 and 87 individuals, where five individuals were capable of theory of mind to one layer of recursion (*d*_1_), and the rest were all naïve (*d*_0_). I simulated 500 instances of each of these populations as described above to 500 iterations, and recorded all trajectories. I then extracted each animal’s Voronoi area, as described above, at every 20th iteration. This was done between *t* = 250 and 350, by which time effects of initial randomisation would not matter any longer.

I calculated separately the mean Voronoi areas of *d*_0_ and *d*_1_ individuals across time in each instance. I then found the proportion of simulations in which the mean areas of *d*_0_ was larger than that of *d*_1_ individuals, and used this proportion as my *d*_1_ survival advantage metric. I then computed the significance of this metric by first quantifying the null distribution, by drawing 5000 permutations of 5 *d*_0_ individuals at a time, and recomputing each time the metric focussing on these 5 individuals. For significance testing, I found the fraction of time the survival advantage metric in these permutations was at least as large as that of true *d*_1_ individuals. This fraction acts as a *p*-value with one tail accounting for stochasticity resulting from finite size effects and initial positions of the animals.

